# Benzo[a]pyrene-induced AHR activation in human ESCs primes premature neurogenesis in brain organoids

**DOI:** 10.64898/2026.04.30.722088

**Authors:** Bohyeon Jeong, Lei Yang, Tharindu Ranathunge, Young-Goo Han

**Affiliations:** Department of Developmental Neurobiology, St. Jude Children’s Research Hospital, 262 Danny Thomas Place, Memphis, TN 38105, USA; Department of Chemical Biology and Therapeutics, St. Jude Children’s Research Hospital, 262 Danny Thomas Place, Memphis, TN 38105, USA

**Keywords:** Polycyclic aromatic hydrocarbons, Benzo[a]pyrene, Aryl hydrocarbon receptor, Neurodevelopment, Premature neurogenesis, Human cerebral organoids

## Abstract

Benzo[a]pyrene (BaP), a representative polycyclic aromatic hydrocarbon (PAH), is a widespread environmental toxicant and potent ligand of the aryl hydrocarbon receptor (AHR). Yet, how early developmental exposure to BaP influences human neurodevelopment remains poorly understood. We first examined AHR expression dynamics during human embryonic stem cell (ESC)-derived cerebral organoid development and found that AHR expression was highest at the ESC stage and declined during subsequent differentiation, suggesting a potential window of heightened susceptibility to AHR-mediated environmental perturbations. Based on this observation, ESCs were exposed to BaP (0.1, 1 μM) for 7 days prior to organoid generation. BaP exposure did not alter proliferation, cell death, or global transcription of ESCs but increased expression of a subset of AHR target genes. Remarkably, however, organoids derived from BaP-exposed ESCs exhibited profound morphological defects resulting from premature neurogenesis, characterized by disrupted neural rosette organization, reduced EOMES⁺ intermediate progenitors, and increased BCL11B⁺ neurons. Pharmacological inhibition of AHR with CH-223191 attenuated AHR activation and rescued the progenitor-neuron imbalance. These findings identify AHR signaling as a critical upstream mediator of BaP-induced developmental neurotoxicity and highlight the vulnerability of early pluripotent stages to environmental insults.

## 1. Introduction

Exposures to environmental pollutants are increasingly recognized as critical contributors to disease etiology, particularly in neurodevelopmental disorders. While genetic factors play a fundamental role, mounting evidence indicates that prenatal exposure to environmental toxicants can durably influence brain development [1–4]. Large-scale biomonitoring studies have revealed widespread exposure to environmental chemicals during pregnancy [1, 5, 6], highlighting the urgency of determining how specific contaminants perturb early human neurodevelopment.

Among airborne toxicants, polycyclic aromatic hydrocarbons (PAHs) are a major class of combustion-derived pollutants of concern. Benzo[a]pyrene (BaP) is a prototypical PAH of particular interest; it is generated through incomplete combustion of organic materials and is ubiquitously detected in traffic-related air pollution, industrial emissions, tobacco smoke, and wildfire-derived particulate matter [7–9]. Reflecting its significance, BaP serves as the primary "surrogate marker" for all atmospheric PAHs for the UK Department for Environment, Food and Rural Affairs and acts as the central reference molecule for the US Environmental Protection Agency’s 16 priority PAHs [10, 11]. BaP is readily adsorbed to fine particulate matter and enters the human body through inhalation, ingestion of contaminated food, and dermal contact. Importantly, BaP and its metabolites cross the placental barrier, resulting in measurable fetal exposure during pregnancy [12–14]. Biomonitoring studies confirming the presence of BaP metabolites and PAH-DNA adducts in maternal blood, cord blood, and placental tissues underscore the direct threat exposed to the fetus during critical developmental windows [12–14].

Epidemiological studies have linked prenatal PAH exposure with adverse neurodevelopmental outcomes, including reduced cognitive performance, behavioral dysregulation, and attention deficits in children [15–17]. Experimental studies in rodent models further demonstrate that gestational BaP exposure disrupts fetal growth and induces persistent neurobehavioral abnormalities in offspring [18, 19]. Despite these associations, the mechanisms by which BaP perturbs early human neurodevelopment remains incompletely understood.

A central pathway mediating BaP toxicity is activation of the aryl hydrocarbon receptor (AHR), a ligand-activated transcription factor that regulates xenobiotic metabolism and diverse developmental programs [20–22]. Upon ligand binding, AHR translocates to the nucleus, dimerizes with aryl hydrocarbon receptor nuclear translocator (ARNT), and induces expression of target genes, including cytochrome P450 enzymes [20, 23, 24]. Beyond detoxification, accumulating evidence indicates that AHR also plays roles in stem cell biology and neural differentiation [21, 25, 26]. Developmental profiling studies have reported high AHR expression in embryonic stem cells (ESCs), which decreases during early differentiation [21, 27]. Genetic disruption of AHR in animal models has also been associated with altered neuronal differentiation and structural abnormalities in the developing nervous system [26, 28–30]. These findings suggest that AHR is not merely a xenobiotic sensor but may actively regulate neurodevelopmental trajectories. Importantly, epidemiological data indicate that prenatal PAH exposure is more consistently associated with neurodevelopmental impairment than postnatal exposure [31], implying that early developmental windows confer heightened susceptibility. However, the specific temporal stages at which BaP exposure exerts the greatest impact and the molecular mechanisms linking early AHR activation to later neurodevelopmental outcomes remain poorly defined in human-relevant systems.

Human pluripotent stem cell-derived cerebral organoids provide a powerful platform to model early cortical development and to interrogate stage-specific vulnerability to environmental toxicants [32–34]. By recapitulating key features of human neurogenesis, including the sequential progression from pluripotent stem cells to neural progenitors and differentiated cortical neurons, this system enables mechanistic dissection of early molecular perturbations and their downstream developmental consequences.

Here, we investigated whether BaP exposure affects cortical neurogenesis using human cerebral organoids. BaP exposure during the ESC stage had minimal immediate effects on ESCs but subsequently disrupted neural progenitor maintenance in cerebral organoids, leading to premature neurogenesis. By integrating transcriptomic profiling with organoid-based phenotypic analyses and pharmacological rescue experiments, we identify AHR as a primary upstream mediator of BaP-induced premature neurogenesis. AHR expression was highest in ESCs and decreased during neural differentiation, suggesting that pluripotent cells have particularly heightened susceptibility to BaP-induced AHR activation with long-lasting effects in subsequent brain development. We further examined whether pharmacological inhibition of AHR signaling mitigates BaP-induced developmental alterations. These findings establish a mechanistic link between early-life BaP exposure, AHR activation, and neurodevelopmental imbalance in a human-relevant model system.

## 2. Materials and Methods

### 2.1. Human ESC Culture and Cerebral Organoid Generation

H9 human ESCs (WiCell Research Institute, Madison, WI, USA) were maintained at 37 °C with 5% CO₂ under feeder-free conditions on Matrigel-coated plates (Corning, Corning, NY, USA) in mTeSR1 medium (STEMCELL Technologies, Vancouver, Canada), with daily medium changes. Cells were passaged every 3-4 days using Versene (Thermo Fisher Scientific, Waltham, MA, USA).

For cerebral organoid generation, ESC colonies were dissociated into single cells using Accutase (Thermo Fisher Scientific) and seeded at 9,000 cells per well in low-attachment v-bottom 96-well plates (MS-9096VZ, S-bio, Hudson, NH) in Essential 8 (E8) medium supplemented with ROCK inhibitor (Y-27632, 50 μM) to generate embryoid bodies (EBs). Neural induction was initiated on day 6 using the DMEM/F12 medium supplemented with N2 (1×), GlutaMAX (1×), MEM non-essential amino acids (1×), and heparin (1 µg/mL). At day 13, organoids were transferred to ultra-low attachment 6-well plates on an orbital shaker (160 rpm) containing differentiation medium composed of a 1:1 mixture of DMEM/F12 and Neurobasal medium supplemented with N2 (0.5×), GlutaMAX (1×), MEM non-essential amino acids (0.5×), antibiotic-antimycotic (1×), β-mercaptoethanol (50 µM), insulin (0.425 µM), and B27 supplement without vitamin A (1×). The WNT activator CHIR99021 (3 µM) was included during early differentiation (days 13-15). From day 16 onward, organoids were maintained in the same differentiation medium without CHIR99021. Beginning on day 25, B27 supplement containing vitamin A was used to promote neuronal maturation and the medium was further supplemented with vitamin C (0.4 mM) and sodium bicarbonate (1 mg/mL). Throughout differentiation, organoids were maintained at 37 °C and 5% CO₂ under continuous orbital shaking (160 rpm) to enhance nutrient diffusion and growth, with medium changes every 2-3 days. All experiments involving human pluripotent stem cells were conducted under approval from the Institutional Review Board (IRB No. 2024-1531-002).

### 2.2. BaP Exposure

Benzo[a]pyrene (BaP; B1760, Sigma-Aldrich, St. Louis, MO, USA) was dissolved in dimethyl sulfoxide (DMSO) to prepare stock solutions. ESCs or differentiating cerebral organoids were exposed to two concentrations (0.1 μM or 1 μM) of BaP to evaluate dose-dependent responses. Exposure concentrations were selected to encompass environmentally relevant ranges reported in human tissues and experimental systems used to model PAH exposure [7, 35–37]. The final DMSO concentration was kept constant across all treatment and control groups.

### 2.3. Intracellular Accumulation of BaP Measurement

#### 2.3.1. Chemicals and Reagents

Acetonitrile (LC-MS grade; Honeywell, Cat# LC015), water (LC-MS grade; Honeywell, Cat# 7732-18-5), and Warfarin (Sigma-Aldrich, Cat# A2250) were used without further purification. Benzo[a]pyrene (BaP) was used as the analyte. All other reagents were of analytical grade.

#### 2.3.2. Sample Preparation

ESCs were dissociated into single cells using Accutase, and cell numbers were counted for normalization. The cells were then pelleted by centrifugation at 300 x g for 5 min and stored at - 80°C for subsequent analysis. Frozen cell pellets were maintained on dry ice and processed in individual 1.5 mL microcentrifuge tubes. Each pellet was quenched with 100 µL of acetonitrile containing 100 µM warfarin as an internal standard. Samples were vortexed (∼15 s) and centrifuged at 13,200 rpm for 2 min (Eppendorf 5415D). Following centrifugation, 80 µL of the supernatant was transferred to a 96-well plate (Corning, Cat# 3363) and mixed with 40 µL of water. The plate was sealed and shaken at 600 rpm for 2 min (IKA MTS 2/4) prior to analysis.

#### 2.3.3. Standard Curve Preparation

A 2 mM stock solution of benzo[a]pyrene (free base equivalent) was prepared in DMSO. Working solutions were generated by serial dilution in acetonitrile containing 100 µM internal standard. A 15 µM solution was prepared by diluting 7.5 µL of stock to 1 mL. Subsequent standards (3, 0.6, 0.12, and 0.024 µM) were prepared by sequential 1:5 dilutions (160 µL into 640 µL diluent; final volume 800 µL). Blank cell pellets were resuspended in 600 µL of water, and 100 µL aliquots were transferred into microcentrifuge tubes. Following centrifugation (13,200 rpm, 2 min) and removal of supernatant, 100 µL of working solution was added to each pellet to generate matrix-matched standards. Standards were processed identically to study cell samples.

#### 2.3.4. Ultra-Performance Liquid Chromatography (UPLC) Conditions

Chromatographic separation was performed on an ACQUITY UPLC BEH C18 1.7 mm, 2.1 x 50 mm column (Waters Corporation, Milford, MA) using an ACQUITY UPLC system. Data acquisition was carried out using MassLynx v4.2 and processed with TargetLynx software. Detection was achieved using a photodiode array (PDA) detector, with the flow subsequently split between an evaporative light scattering detector (ELSD) and a single quadrupole mass spectrometer (SQD2). The mobile phases consisted of water containing 0.1% formic acid (A) and acetonitrile containing 0.1% formic acid (B). The injection volume was 10 µL. The column temperature was maintained at 63 °C, and the sample temperature at 20 °C. The gradient was run at a flow rate of 1.0 mL/min as follows: 0–0.2 min, 70% A; 0.2–1.4 min, linear gradient to 5% A; 1.4–1.95 min, held at 5% A; and 1.95–2.0 min, returned to 70% A [38].

### 2.4. BrdU incorporation assay

To assess cell proliferation, ESCs were exposed to each chemical for 5 days, followed by the addition of 10 μM 5-bromo-2’-deoxyuridine (BrdU) for overnight incubation. The following day, the medium was replaced, and cells were cultured until day 7. Cells were then fixed with 4% paraformaldehyde (PFA) for 15 min at room temperature. Following fixation, cells were permeabilized with 0.3% Triton X-100 for 20 min and treated with 1.5 N HCl for 30 min at room temperature to denature DNA. Acid was neutralized using 0.1 M Tris buffer (pH 8.8) for 20 min. After blocking with 3% bovine serum albumin (BSA), cells were incubated with sheep anti-BrdU (ab1892, 1:250) overnight at 4°C, followed by incubation with Alexa Fluor^tm^ 568-conjugated secondary antibody (Thermo Fisher Scientific). Nuclei were counterstained with DAPI. Fluorescence images were acquired using an Axio Imager microscope (Zeiss, Oberkochen, Germany) equipped with an Apotome optical sectioning system.

### 2.5. Quantitative Real-Time PCR (qRT-PCR)

At designated differentiation stages, cells or organoids were harvested, and total RNA was extracted using TRIzol Reagent (Invitrogen, Carlsbad, CA, USA). RNA was purified using the RNA Clean & Concentrator™-5 kit with DNase I treatment (Zymo Research, Irvine, CA, USA). Complementary DNA (cDNA) was synthesized using SuperScript™ VILO™ Master Mix (Invitrogen). Quantitative PCR was performed using PowerUp™ SYBR™ Green Master Mix (Applied Biosystems, Carlsbad, CA, USA) on a QuantStudio™ 7 Flex Real-Time PCR System (Applied Biosystems). Gene expression levels were normalized to *ACTB* and calculated using the comparative ΔΔCt method. Primer sequences were as follows:

- OCT4 (F 5′-CCCGAAAGAGAAAGCGAACC-3′; R 5′-TACAGAACCACACTCGGACC-3′)
- SOX2 (F 5′-CGGAAAACCAAGACGCTCAT-3′; R 5′-TTCATGTGCGCGTAACTGTC-3′)
- NANOG (F 5′-TGTCTTCTGCTGAGATGCCT-3′; R 5′-TTTCTTGACCGGGACCTTGT-3′)
- AHR (F 5′-TTGAACCATCCCCATACCCC-3′; R 5′-TTCTGGCTGGCACTGATACA-3′)
- AHRR (F 5′- GAGACTCCAGGACCCACAAA-3′; R 5′-AGATGGGCGAGTTCCTGAAA-3′)
- CYP1A1 (F 5′-AGGTCCTTAGGCCTCTGAGA-3′; R 5′-GGCTGTCTCTTCCCTTCACT-3′)
- CYP1B1 (F 5′-GAAACCTCGACTTTGCCAGG-3′; R 5′-CAAGACGTCAACAGGAACCC-3′)
- ACTB (F 5′-GCTCACCATGGATGATGATATCGC-3′; R 5′-CACATAGGAATCCTTCTGACCCAT-3′)

### 2.6. Lactate Dehydrogenase (LDH) Cytotoxicity Assay

Cytotoxicity was assessed using a Lactate Dehydrogenase (LDH) Cytotoxicity Assay Kit (HY-K1090, MedChemExpress, Monmouth Junction, NJ, USA) according to the manufacturer’s instructions. Culture supernatants were collected and incubated with LDH reaction mixture. For positive control samples, cells were treated with the provided lysis buffer to determine maximum LDH release. Absorbance was measured at 490 nm using a microplate reader (Epoch Microplate Spectrophotometer, BioTek, Winooski, VT, USA). Cytotoxicity was calculated as a percentage of maximum LDH release.

### 2.7. Brightfield Imaging and Morphological Analysis

Organoids were imaged at indicated time points using a CKX53 inverted microscope (Olympus, Tokyo, Japan). Organoid size and morphology were analyzed using ImageJ software (NIH, Bethesda, MD, USA). Diameters were measured from brightfield images and quantified for each experimental group.

### 2.8. Immunofluorescence Staining

Cells or organoids were fixed in 4% PFA at 4°C for 1 h. Organoids were cryoprotected in 30% sucrose overnight, embedded in OCT compounds (Fisher Healthcare) containing 4% sucrose, and cryosectioned at 16 μm thickness. Samples were permeabilized and blocked in blocking buffer (3% bovine serum albumin in PBS) before incubation with primary antibodies overnight at 4°C. We used the following primary antibodies: SOX2 (sc17320, 1:500), CDH2 (alias N-cadherin; 22018-1-AP, 1:500), phospho-histone H3 (PHH3; 06-570, 1:200), MKI67 (alias Ki67; M7240, 1:200), CDKN1A (alias p21; sc6246, 1:100), Cleaved Caspase-3 (9661S, 1:400), EOMES (alias TBR2; 14-4877-82, 1:200), BCL11B (alias CTIP2; ab18465, 1:500), S100B (AMAb91038, 1:500), and RBFOX3 (alias NEUN; GTX132974, 1:500). Following incubation with Alexa Fluor^tm^-conjugated secondary antibodies (Thermo Fisher Scientific), nuclei were counterstained with DAPI. Images were acquired using an Axio Imager fluorescence microscope (Zeiss, Oberkochen, Germany) with the Apotome optical sectioning system.

### 2.9. RNA sequencing and analysis

Total RNA was extracted from ESCs using TRIzol and purified as described above. A sequencing library was prepared using a TruSeq Stranded Total RNA Kit (Illumina) and sequenced with an Illumina HiSeq system. Total stranded RNA sequencing data were processed by the internal AutoMapper pipeline. Briefly the raw reads were first trimmed (Trim-Galore version 0.60), mapped to human (GRCh38) [39] and then the gene level values were quantified (RSEM v1.31) [40] based on GENCODE annotation (v31 for human, VM22 for mouse). Genes with low counts (counts per million [CPM] < 0.1) were removed from the analysis, and only protein-coding genes were used for differential expression analysis. Normalization factors were generated using the TMM method [41]. Counts were transformed using voom [42] and analyzed using the lmFit and eBayes functions (R limma package version 3.42.2) [43]. The false discovery rate (FDR) was estimated using the Benjamini–Hochberg method. Pre-ranked Gene set enrichment analysis (GSEA) [44] was carried out using negative log10 (P.Value)*log2FoldChange value (from differential expression analysis) ranked gene list against gene sets in the Molecular Signatures Database (MSigDB v2023.1). GSEA version 4.3.2 was used with following parameters: number of permutations =1000, permutation type = gene_set, metric for ranking genes = Signal2Noise, Enrichment statistic = Weighted.

### 2.10. CH-223191 Pretreatment

To inhibit BaP-induced AHR activation, ESCs were treated with CH-223191 (10 μM; 72732, STEMCELL Technologies, Vancouver, Canada), a selective competitive AHR antagonist, 1 h prior to BaP exposure [45, 46]. During the 1-week exposure period, CH-223191 pretreatment and BaP treatment were repeated daily at each medium change (**Fig. 7B**).

### 2.11. Cytosolic and Nuclear Fractionation and Western Immunoblotting

Cytoplasmic and nuclear extracts were prepared using NE-PER™ Nuclear and Cytoplasmic Extraction Reagents (Thermo Fisher Scientific) according to the manufacturer’s protocol. Equal amounts of protein (25 μg) were separated on 6-18% gradient SDS-PAGE gels and transferred to PVDF membranes. Membranes were incubated with primary antibodies against AHR (GTX129013, GeneTex, Irvine, CA), GAPDH (60004-1-Ig, Proteintech, Rosemont, IL), and Lamin B1 (ab16048, Abcam, Waltham, MA). Protein bands were visualized using chemiluminescence with a ChemiDoc Go Imaging System (Bio-Rad) and quantified using ImageJ software. AHR levels were normalized to GAPDH (cytosolic marker) or Lamin B1 (nuclear marker). Nuclear-to-cytosolic AHR ratios were then calculated for each sample.

### 2.12. Image Analysis

Fluorescence images were analyzed using ImageJ software (National Institutes of Health). For quantification of marker-positive cells, images from at least three independent experiments were analyzed. The number of positive cells was manually counted or quantified using threshold-based analysis and normalized to the total number of DAPI-stained nuclei. All image processing and quantification were performed under identical settings across experimental groups.

### 2.13. Statistical Analysis

All experiments were performed with at least three independent biological replicates. Data were analyzed using one-way ANOVA followed by Dunnett’s or Tukey’s multiple comparisons test, as appropriate. Statistical analyses were performed using GraphPad Prism. A *p*-value < 0.05 was considered statistically significant. Data are presented as mean ± the standard error of the mean (SEM).

## 3. Results

### 3.1. ESCs express high levels of AHR and accumulate BaP while maintaining pluripotency, proliferation, and viability

Since PAHs are known to exert their biological effects through AHR signaling [20, 47], we first examined the dynamic expression of *AHR* during brain organoid development under our culture conditions (**Fig. 1A**). qPCR analysis across developmental stages, from ESCs to embryoid bodies (EB), neural induction (NI), early differentiation (ED), and maturation (Mat), revealed that *AHR* mRNA was highest at the ESC stage and sharply declined following EB formation. A modest level of *AHR* was observed during early differentiation, followed by a decline at the maturation stage. Based on this mRNA expression pattern, we focused on ESC-stage exposure, when *AHR* levels were highest.

**Figure 1.**
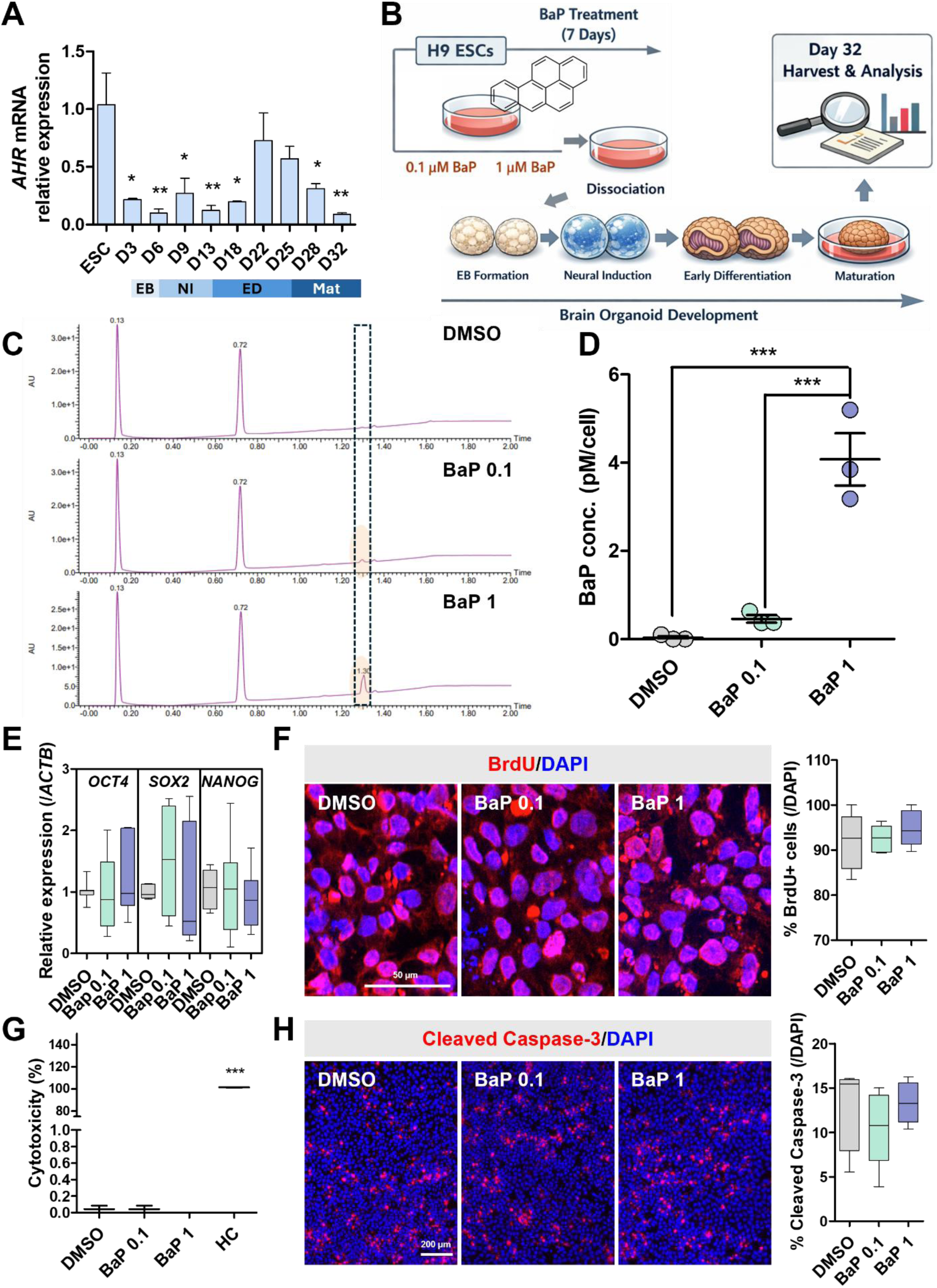
ESCs express high levels of AHR and accumulate BaP while maintaining pluripotency, proliferation, and viability. **(A)** Relative expression levels of *AHR* during organoid development from the ESC stage through embryoid body formation (EB), neural induction (NI), early differentiation (ED), and maturation (Mat). **(B)** Schematic diagram illustrating the experimental design. Human ESCs were exposed to BaP for 7 days prior to organoid generation, followed by organoid formation and downstream analyses. **(C)** Representative UPLC-MS chromatograms showing BaP detection in ESC lysates after 7-day exposure. The BaP-specific peak at retention time of 1.3 min is indicated by a dotted box. **(D)** Intracellular accumulation of BaP in ESC after 7-day exposure. **(E)** Box plots showing qPCR analysis of pluripotency markers (*OCT4, SOX2*, and *NANOG*) in ESCs after 7 days of exposure under each condition. **(F)** Representative immunofluorescence images and quantifications of BrdU^+^ ESCs. **(G)** LDH assay results assessing cytotoxicity following 7-day exposure. HC: high control (lysis buffer-treated, 100% LDH release). **(H)** Representative immunofluorescence images and quantifications of Cleaved Caspase-3^+^ ESCs (red). Values denote mean ± SEM. *\*p<0.05, **p<0.005, ***p<0.0005*. Scale bars: **(F)** 50 μm, **(H)** 200 μm.

Human ESCs were exposed to BaP for 7 days prior to organoid generation, followed by morphological and cellular analyses of the resulting day 32 organoids (**Fig. 1B**). To confirm intracellular accumulation of BaP, ESCs were exposed to BaP for 7 days and analyzed by UPLC-MS. A peak corresponding to BaP was detected at a retention time of approximately 1.3 min, and its signal intensity increased in a dose-dependent manner (**Fig. 1C** and **D**), supporting intracellular accumulation of BaP following exposure. To determine whether BaP exposure affects ESC identity, we assessed expression levels of the pluripotency markers, OCT4, SOX2, and NANOG, by qPCR (**Fig. 1E**). BaP did not alter transcript levels of these markers. Furthermore, RNA sequencing revealed no substantial changes in global gene expression; principal component analysis (PCA) and hierarchical clustering analysis failed to separate control and BaP-treated cells (**Fig. 5**, detailed in a later section), indicating that BaP exposure did not disrupt ESC identity. Consistent with no change in global gene expression, BaP did not alter the number of BrdU-incorporating proliferating ESCs (**Fig. 1F**). Lastly, we evaluated cytotoxicity. LDH assays showed no significant increase in LDH release compared to DMSO controls (**Fig. 1G**), suggesting the absence of necrotic cell death. Immunostaining for cleaved caspase-3 also revealed no significant differences in the number of apoptotic cells between groups (**Fig. 1H**). Together, these results indicate that BaP exposure did not induce overt changes in identity, proliferation, or death of ESCs.

### 3.2. ESC-stage BaP exposure, but not later exposure, disrupts cerebral organoid morphology

We generated organoids using ESCs pretreated with either BaP or DMSO for 7 days and monitored their morphology throughout development (**Fig. 2A, B**). BaP pretreatment did not affect early organoid development; however, compared with DMSO-pretreated controls, it caused a progressive increase in organoid diameter (normalized to day 3 EB diameter), particularly at later stages (**Fig. 2B**). At day 32, most BaP-derived organoids displayed prominent irregular surface protrusions (**Fig. 2C**, red arrows), which were rarely found in control-derived organoids. These structural abnormalities were not accompanied by early ESC alterations (**Fig. 1**), suggesting that BaP exposure during the pluripotent stage alters subsequent developmental program rather than inducing acute cytotoxicity.

**Figure 2.**
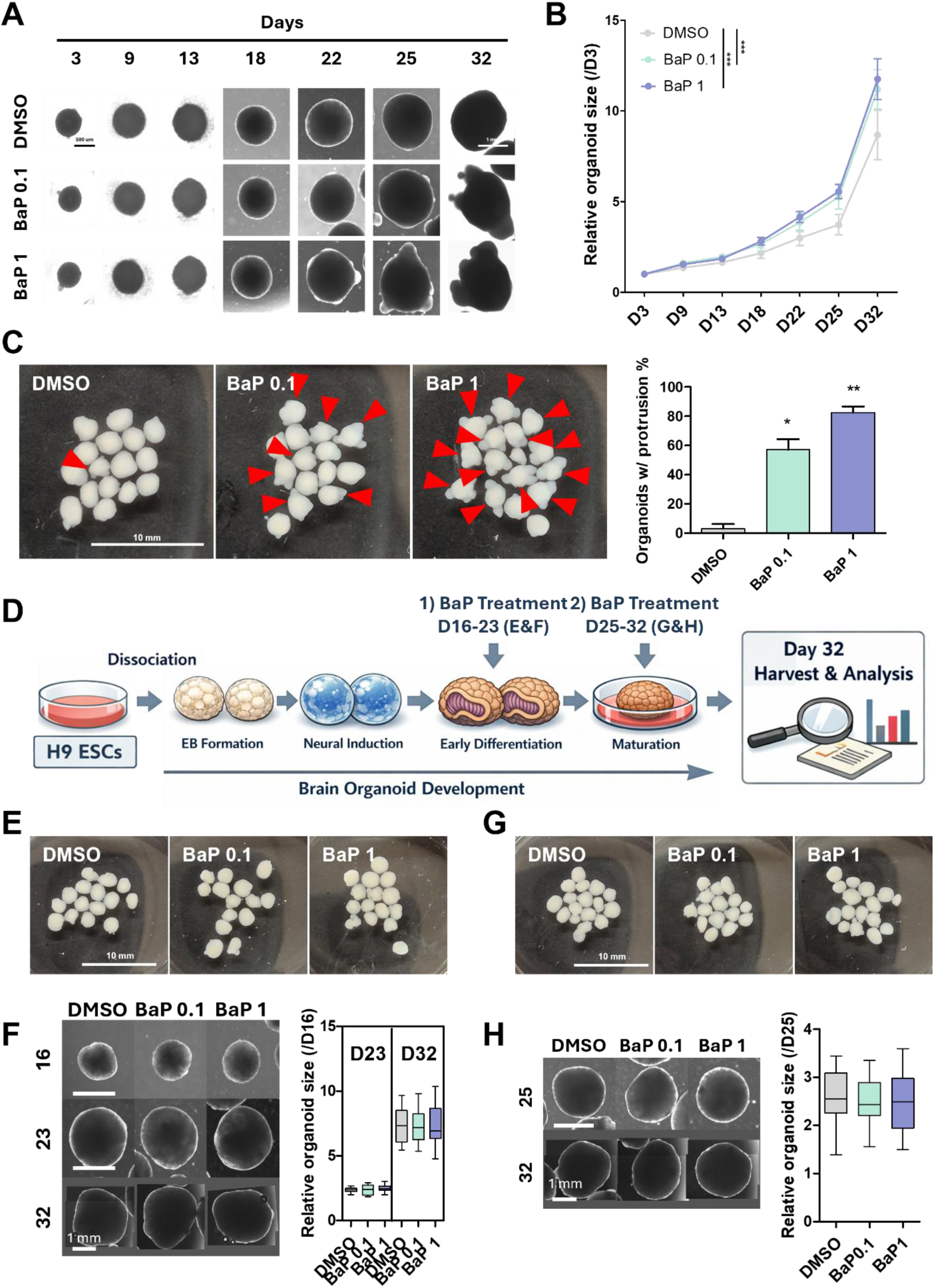
BaP exposure during the ESC stage alters organoid morphology. **(A)** Representative bright-field images showing organoid morphology over time across experimental groups. **(B)** Line graph showing organoid size normalized to day 3, when EBs formed. Organoids derived from BaP-exposed ESCs (BaP 0.1 µM and BaP 1 µM) exhibit increased relative size over time compared with DMSO controls. **(C)** Representative images of organoid morphology at day 32. BaP-treated groups display increased formation of protrusions (red arrows). The accompanying graph shows a higher percentage of organoids with protrusions in BaP-derived groups compared to controls. **(D)** Schematic diagram illustrating experimental design. BaP was applied for 1 week during either 1) early differentiation (day 16-23; corresponding to panels **E** and **F**) or 2) maturation (day 25-32; corresponding to panels **G** and **H**). **(E)** Representative bright-field images of day 32 organoids exposed to DMSO or BaP for 1 week during early differentiation (day 16-23). **(F)** Representative images showing organoid morphology over time during early differentiation exposure, with the accompanying graph indicating relative organoid size at each time point normalized to day 16. **(G)** Representative bright-field images of day 32 organoids exposed to DMSO or BaP for 1 week during maturation (day 25-32). **(H)** Representative images showing organoid morphology over time during maturation exposure, with the accompanying graph indicating relative organoid size at day 32 normalized to day 25. Values denote mean ± SEM. *\*p<0.05, **p<0.005, ***p<0.0005*. Scale bars: **(A)** 500 μm (days 3-25) and 1 mm (days 32), **(C)** 10 mm **(E, G)** 10 mm, **(F, H)** 1 mm.

We next examined whether BaP exposure during organoid development would produce morphological changes similar to those observed following ESC-stage exposure. To address this, organoids were exposed to BaP for one week during either the early differentiation stage (day 16-23) or the maturation stage (day 25-32) (**Fig. 2D**), when AHR is modestly expressed (**Fig. 1A**). BaP exposure during these stages caused no obvious morphological changes in day 32 organoids (**Fig. 2E** and **G**). Additionally, longitudinal imaging throughout the exposure period showed comparable growth trajectories among all groups (**Fig. 2F** and **H**). Collectively, these findings indicate that susceptibility to BaP is highly dependent on developmental timing, with exposure during the ESC stage, when AHR expression is highest, leading to abnormal structural growth of cerebral organoids, whereas exposure during organoid stage does not affect gross morphology or growth dynamics. These results suggest that the ESC stage is a particularly vulnerable window for BaP-induced neurodevelopmental disruption.

### 3.3. BaP exposure during the ESC stage disrupts neural rosette organization and reduces proliferative capacity in cerebral organoids

To further characterize the structural abnormalities observed in organoids derived from BaP-exposed ESCs, we examined internal cytoarchitecture at day 32. Immunostaining for SOX2 and N-cadherin revealed that SOX2-expressing neural progenitors form well-organized neural rosettes along the periphery of the control organoids. Strikingly, neural rosettes were absent in many regions of the BaP-derived organoids, especially in the irregular protrusions (**Fig. 3A**). Irregular protrusions occupied a larger proportion of organoid cross-sectional areas in BaP-derived groups (**Fig. 3B**). Consistent with this, BaP exposure markedly reduced the total number of neural rosettes per organoid (**Fig. 3C**). Furthermore, individual rosette size was also significantly decreased (**Fig. 3D**), with a shift in size distribution toward smaller rosettes in BaP-derived organoids (**Fig. 3E**).

**Figure 3.**
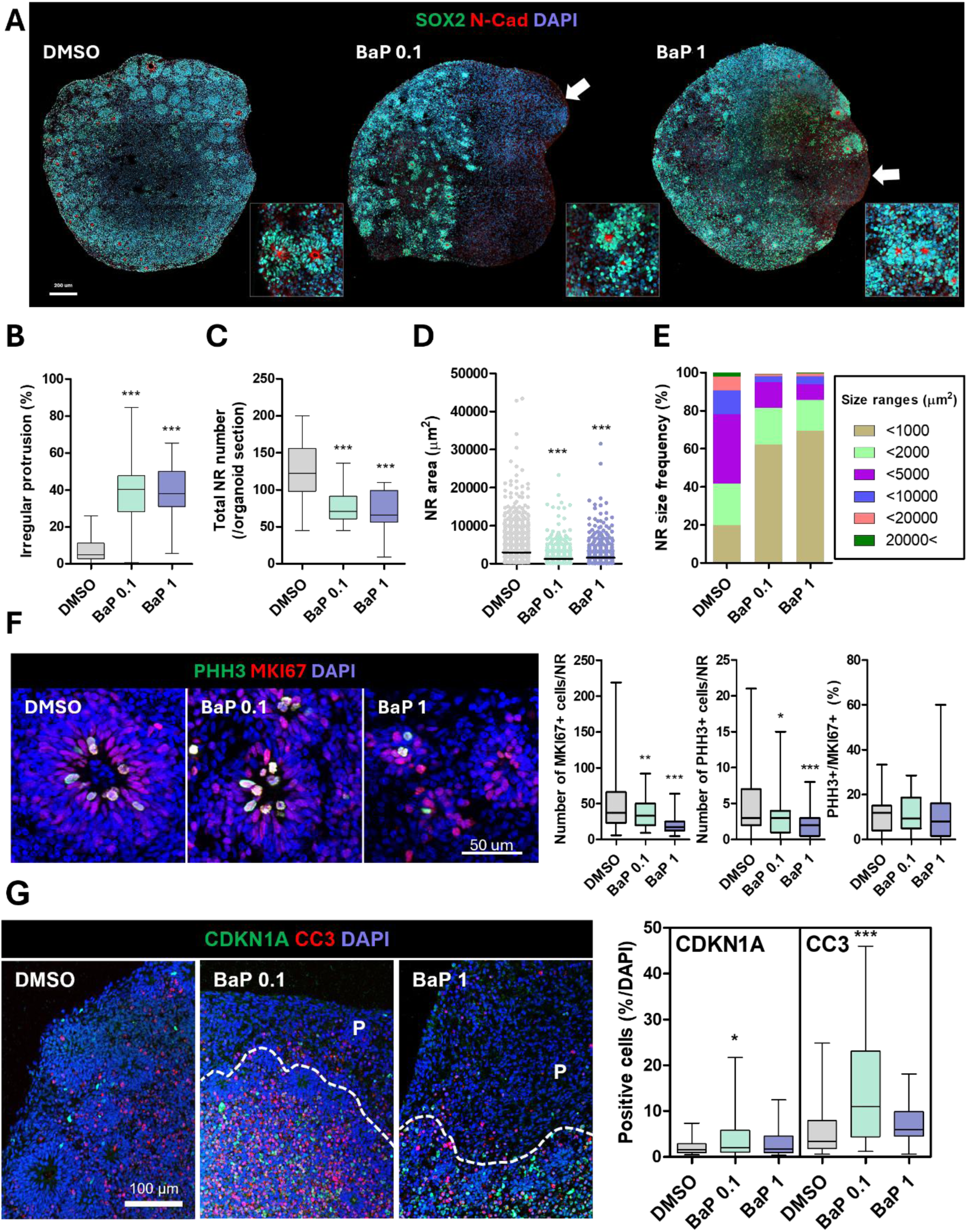
BaP exposure during the ESC stage alters neural rosette organization and proliferative activity in organoids. **(A)** Representative immunofluorescence images of day 32 organoids stained for SOX2 (green), N-Cadherin (red), and DAPI (blue). Insets show magnified views of neural rosettes from each group. White arrows indicate protrusion areas lacking neural rosettes. **(B)** Box plot showing the percentage of organoid cross-sectional area occupied by irregular protrusions. **(C)** Box plot showing the number of neural rosettes (NR) per organoid section. **(D)** Dot plot showing individual neural rosette (NR) size in organoids. **(E)** Graph illustrating the frequency distribution (%) of rosette sizes across experimental groups. **(F)** Representative immunofluorescence images stained for PHH3 (green), MKI67 (red), and DAPI (blue). Box plots show the number of MKI67⁺ cells and PHH3⁺ cells per rosette, as well as the percentage of PHH3⁺ cells among MKI67⁺ cells (PHH3⁺/MKI67⁺, %). **(G)** Representative immunofluorescence images of day 32 organoids stained for CDKN1A (green), Cleaved Caspase-3 (CC3; red), and DAPI (blue), with corresponding box plots showing the percentage of CDKN1A⁺ and CC3⁺ cells. P indicates protrusion; dotted lines demarcate the boundary between protrusion regions and adjacent rosette areas. Values denote mean ± SEM. *\*p < 0.05, **p < 0.005, ***p < 0.0005*. Scale bars: **(A)** 200 μm**, (F)** 50 μm, **(G)** 100 μm.

Given the reduction in rosette number and size, we next assessed proliferative activity. Immunostaining for MKI67 and phospho-histone H3 (PHH3) demonstrated a significant decrease in the number of MKI67⁺ and PHH3⁺ cells per rosette in BaP-treated organoids (**Fig. 3F**). However, the proportion of PHH3⁺ cells among MKI67⁺ cells (PHH3⁺/MKI67⁺, %) was not significantly altered, indicating that while the total proliferative cell pool was reduced, cell-cycle progression within the remaining proliferating population was largely preserved.

To determine whether additional mechanisms contributed to rosette shrinkage, we examined markers of cell-cycle arrest and apoptosis. Organoids generated from ESC treated with 0.1 µM of BaP showed a significant increase in the number of cells expressing CDKN1A, a cyclin-dependent kinase inhibitor and a marker of cell cycle arrest, near neural rosette regions, whereas the 1 µM-derived organoids displayed only a modest, non-significant increase (**Fig. 3G**). Similarly, cleaved caspase-3, an apoptosis marker, levels were significantly elevated at 0.1 µM but showed only a slight upward trend at 1 µM without reaching statistical significance. Together, these findings suggest that BaP exposure reduced the neural progenitor pool primarily through impaired proliferative capacity, whereas cell-cycle arrest and apoptosis appear to contribute only to a limited extent to the observed rosette shrinkage.

### 3.4. BaP exposure promotes premature neuronal differentiation and alters neural lineage composition

We next investigated whether reduced rosette size and progenitor number were associated with altered lineage specification. Immunostaining for SOX2, EOMES (alias TBR2), and BCL11B (alias CTIP2) revealed significant changes in neural lineage composition (**Fig. 4A**). BaP decreased the proportion of EOMES⁺ intermediate progenitor cells with a concomitant increase in BCL11B⁺ early born neurons relative to SOX2⁺ neural stem cells in organoids (**Fig. 4B**). Low-magnification imaging further showed that protrusive regions lacking organized rosette structures were predominantly composed of BCL11B⁺ neurons (**Fig. 4C**). Most cells in the protrusion area also expressed RBFOX3 (alias NEUN), another neuronal marker (**Fig. 4D**). These results suggest that the abnormal outgrowth reflects premature neuronal differentiation rather than simple structural deformation. To assess glial differentiation, we stained for S100B (**Fig. 4D**). The number of S100B⁺ astroglial cells also slightly increased in BaP-treated organoids. Collectively, these results indicate that BaP exposure at the ESC stage leads to reduced progenitor maintenance and a shift toward premature neuronal and glial differentiation, ultimately resulting in disrupted rosette architecture and protrusions filled with prematurely differentiated neurons in cerebral organoids.

**Figure 4.**
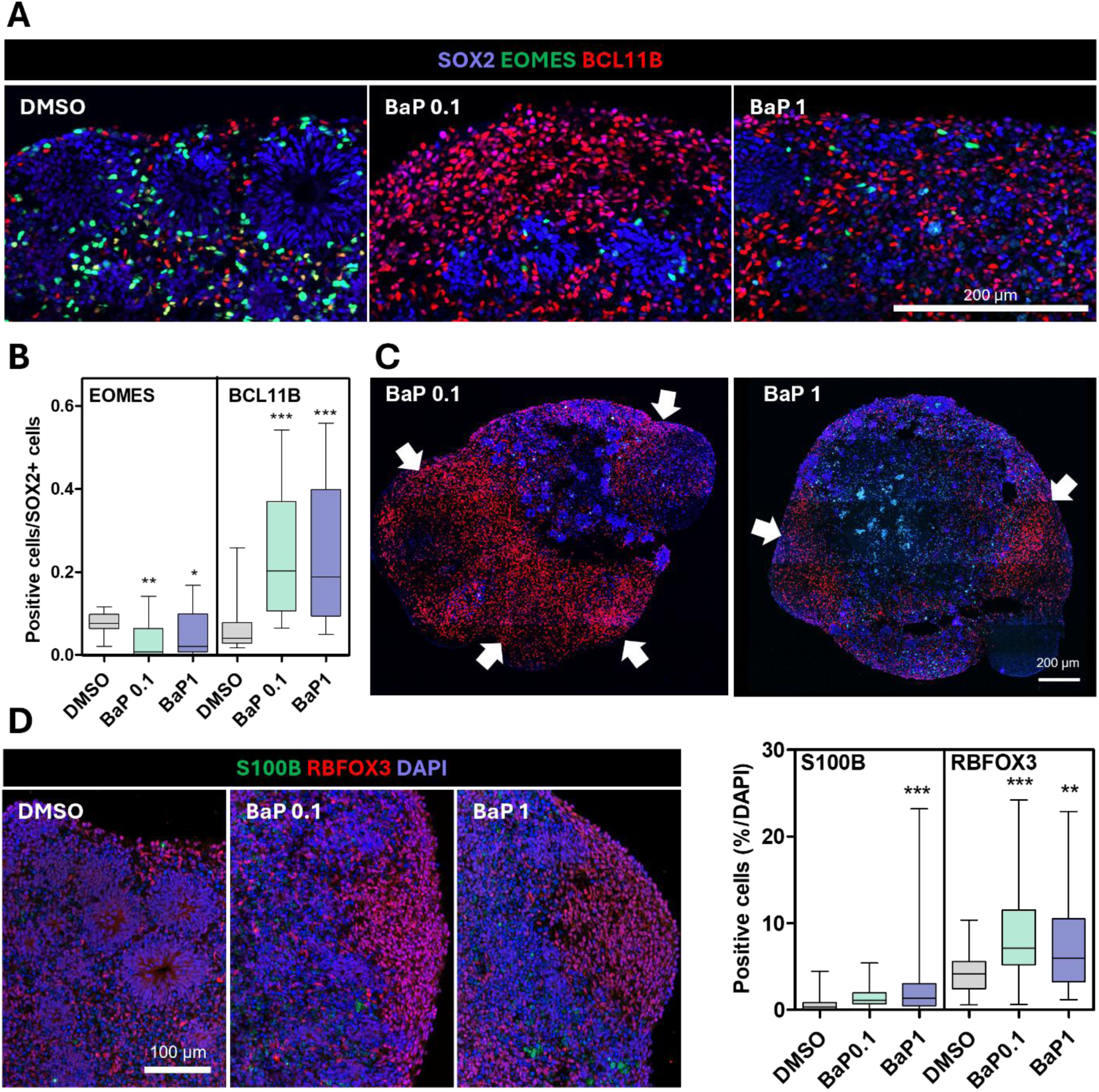
BaP exposure during the ESC stage alters neural lineage composition in organoids. **(A)** Representative immunofluorescence images of SOX2 (blue), EOMES (green), and BCL11B (red). **(B)** Box plots showing the proportion of EOMES⁺ intermediate progenitor cells and BCL11B⁺ neurons relative to SOX2⁺ cells in organoids. **(C)** Low-magnification images of BaP organoids with protrusions (white arrows), which are predominantly composed of BCL11B⁺ neurons. **(D)** Representative immunofluorescence images of S100B (green), RBFOX3 (red), and DAPI (blue), with box plots showing the percentage of S100B⁺ glial cells and RBFOX3⁺ neurons. Values denote mean ± SEM. *\*p < 0.05, **p < 0.005, ***p < 0.0005*. Scale bars: **(A)** 200 μm, **(C)** 200 μm, **(D)** 100 μm.

### 3.5. BaP exposure induces AHR-associated transcriptional reprogramming in ESCs

To investigate early molecular alterations induced by BaP exposure in pluripotent cells, we performed bulk transcriptomic profiling of ESCs treated with 1 µM BaP. Unsupervised analyses, including PCA and hierarchical clustering, did not reveal clear segregation of samples by treatment condition, with samples distributed without distinct grouping patterns (**Fig. 5A, B**). These results suggest that BaP exposure does not induce large-scale global transcriptional shifts at the ESC stage. Differential expression analysis identified 86 genes at nominal significance (*p* < 0.05), among which only two genes, *AC008878.3* and *LRAT*, remained significant after multiple-testing correction (*FDR* < 0.0005) (**Fig. 5C**). This limited DEG number was consistent with the preserved ESC pluripotency without changes in proliferation or cell death following BaP treatment (**Fig. 1**).

**Figure 5.**
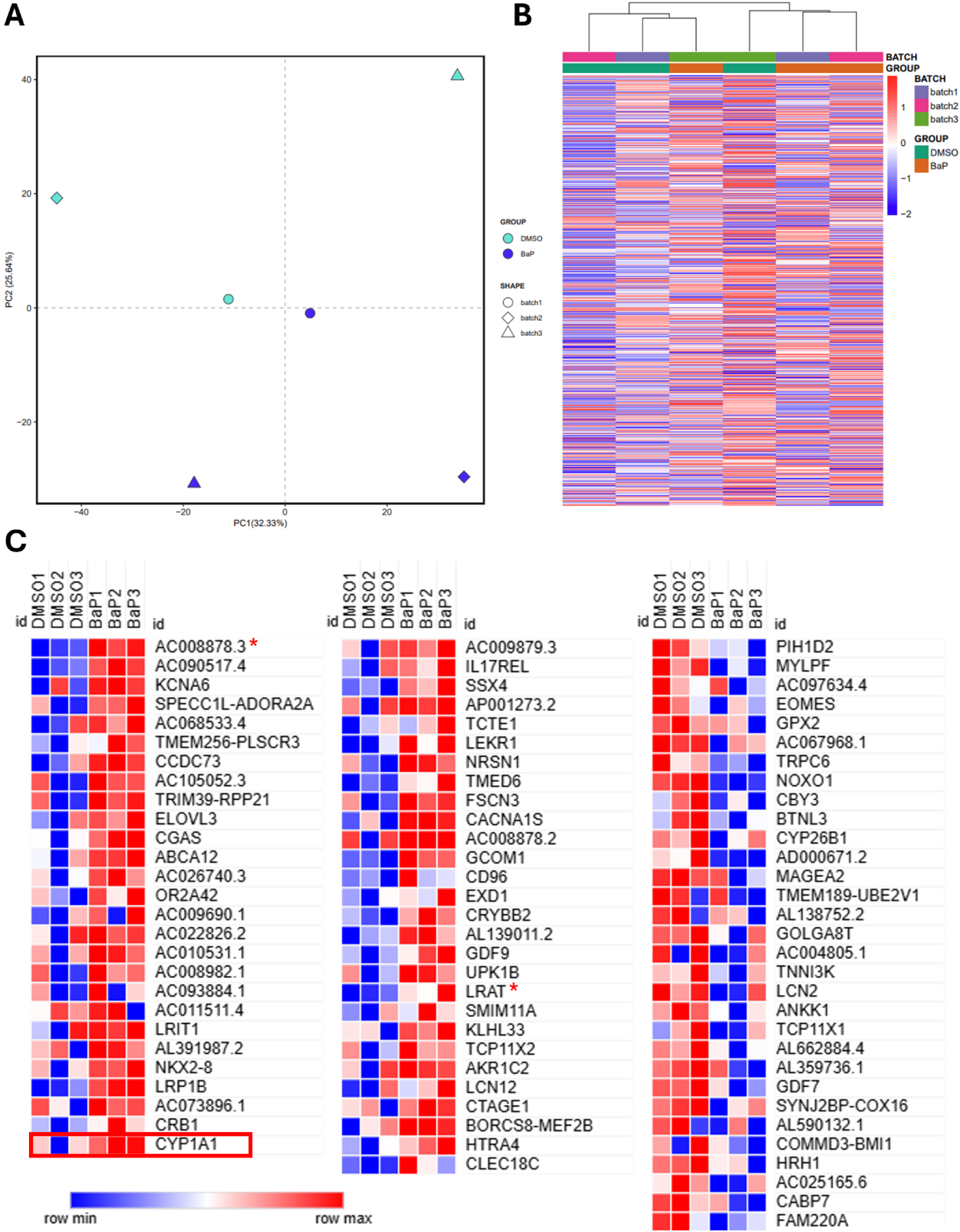
Bulk transcriptomic profiling reveals limited transcriptional changes in BaP-exposed ESCs. **(A)** Principal component analysis (PCA) and **(B)** Hierarchical clustering of transcriptomes of ESC samples treated with 1 µM BaP or DMSO control for 7 days. **(C)** Heat map showing 86 candidate differentially expressed genes (nominal *p* < 0.05) between BaP-treated and control ESCs. Genes maintaining significance after False Discovery Rate correction (FDR < 0.05) are indicated by asterisks (*).

GO enrichment analysis (Biological Process) of 86 differentially expressed genes with nominal significance demonstrated significant enrichment of piRNA processing, cellular response to copper ion, response to copper ion, olefinic compound metabolic process, and estrogen metabolic process (**Fig. 6A**). Notably, protein-protein interaction (PPI) network analysis revealed that *CYP1A1* functioned as a central node linking four of the five enriched processes (all except piRNA processing), suggesting that xenobiotic metabolism-related transcriptional regulation represents a key convergent mechanism in BaP-exposed ESCs. Consistently, KEGG pathway analysis identified chemical carcinogenesis-DNA adducts pathway as significantly enriched (**Fig. 6B**). Although chemical carcinogenesis pathways have been widely reported in differentiated cells, both in culture and in *in vivo* BaP exposure models [48–52], their enrichment in ESCs indicates that canonical xenobiotic response programs are already inducible at the pluripotent stage. Within the KEGG network, *CYP1A1* and *CYP2C8* emerged as core factors, further supporting the activation of cytochrome P450-mediated detoxification machinery. Gene set enrichment analysis (GSEA) reinforced these findings by demonstrating significant enrichment of AHR signaling and cytochrome P450-related gene sets in BaP-treated ESCs (**Fig. 6C**). Consistent with the activation of this pathway at the transcriptional level, qPCR analysis confirmed a robust upregulation of the canonical AHR target gene *CYP1A1* following BaP exposure (**Fig. 6D**).

**Figure 6.**
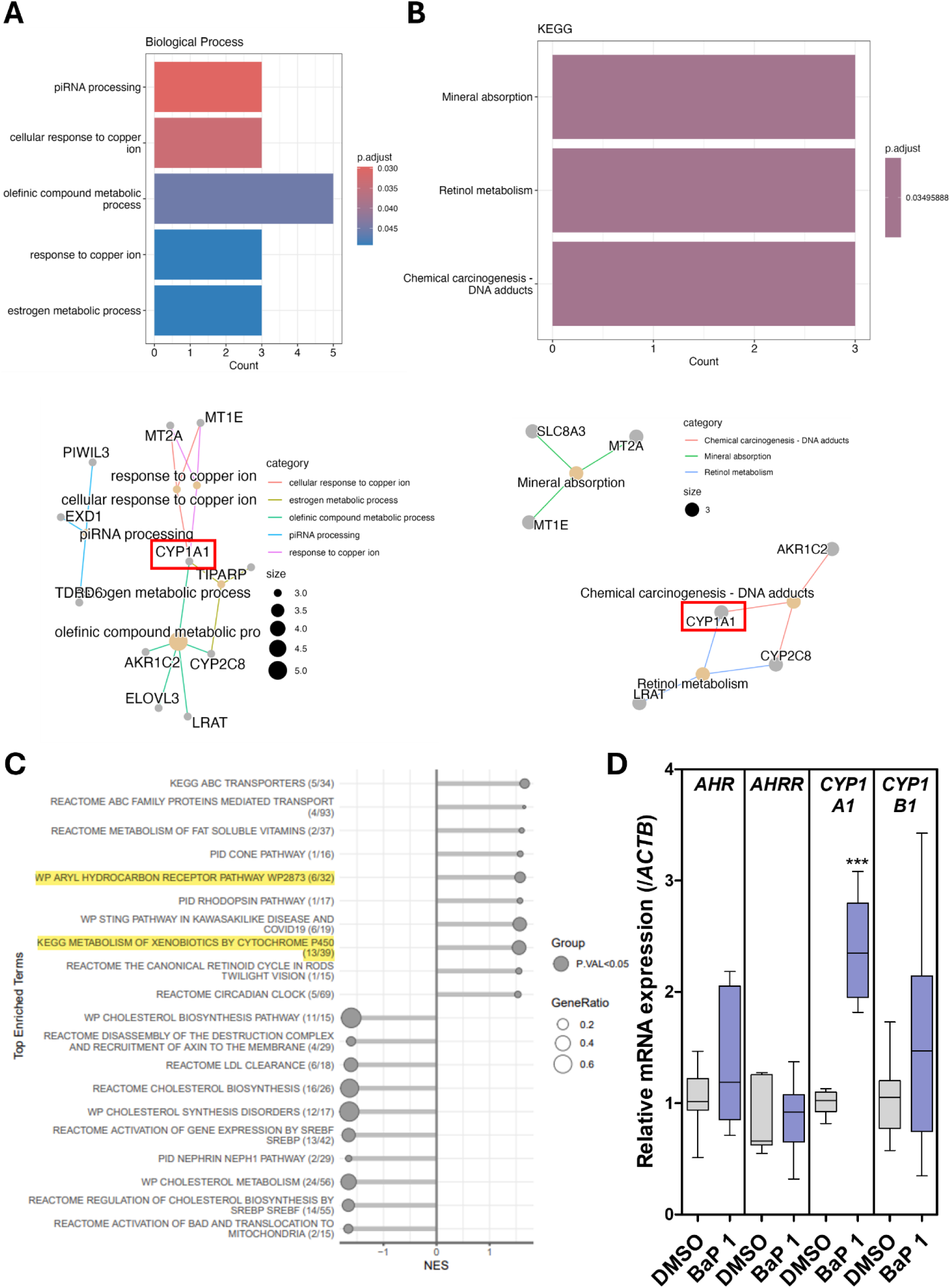
BaP exposure activates AHR-related signaling in ESCs. **(A)** GO enrichment analysis (Biological Process) bar and protein-protein interaction (PPI) network plot and **(B)** KEGG pathway analysis bar and PPI network plot of 86 candidate differentially expressed genes in ESCs exposed to 1 µM BaP. **(C)** GSEA bubble plots showing enrichment of AHR signaling and cytochrome P450-related pathways in BaP-exposed ESCs. **(D)** qPCR results showing expression of the AHR target genes in ESCs following 1 µM BaP exposure. Values denote mean ± SEM. *\*\*\*p < 0.0005*.

Beyond canonical xenobiotic signaling, transcriptomic analyses also identified enrichment of additional metabolic and transport-related pathways following BaP exposure. GSEA showed upregulation of ABC transporter-related pathways, including KEGG ABC transporters and Reactome ABC family protein-mediated transport, consistent with activation of cellular efflux programs involved in xenobiotic export [53, 54]. Retinoid-associated pathways, including metabolism of fat-soluble vitamins and retinol metabolism, were also enriched, whereas cholesterol biosynthesis-related pathways, such as cholesterol biosynthesis, cholesterol metabolism, and SREBP-regulated gene sets, were downregulated. These pathway-level changes indicate that BaP exposure alters multiple metabolic programs in ESCs in parallel with AHR-responsive transcriptional activity. Importantly, despite the absence of pronounced global transcriptional disruption or cytotoxic phenotypes, these coordinated changes indicate that BaP exposure activated AHR-dependent signaling and adaptive metabolic responses in ESCs. The relatively modest DEG quantity may explain the maintenance of pluripotent morphology and viability at this stage. However, such early transcriptional priming could persist and influence downstream lineage specification during later stages of differentiation. Together, these findings suggest that early BaP-induced activation of AHR signaling in ESCs may prime transcriptional programs that influence subsequent neurodevelopmental trajectories during organoid differentiation.

### 3.6. AHR antagonism rescues BaP-induced AHR activation and aberrant neurogenic differentiation

We therefore next used CH-223191, a potent and selective competitive antagonist of the AHR [45, 46], to examine whether pharmacological inhibition of AHR signaling could rescue BaP-induced alterations. CH-223191 blocks ligand-induced AHR nuclear translocation and subsequent DNA binding, thereby preventing transcriptional activation of AHR target genes (**Fig. 7A**).

**Figure 7.**
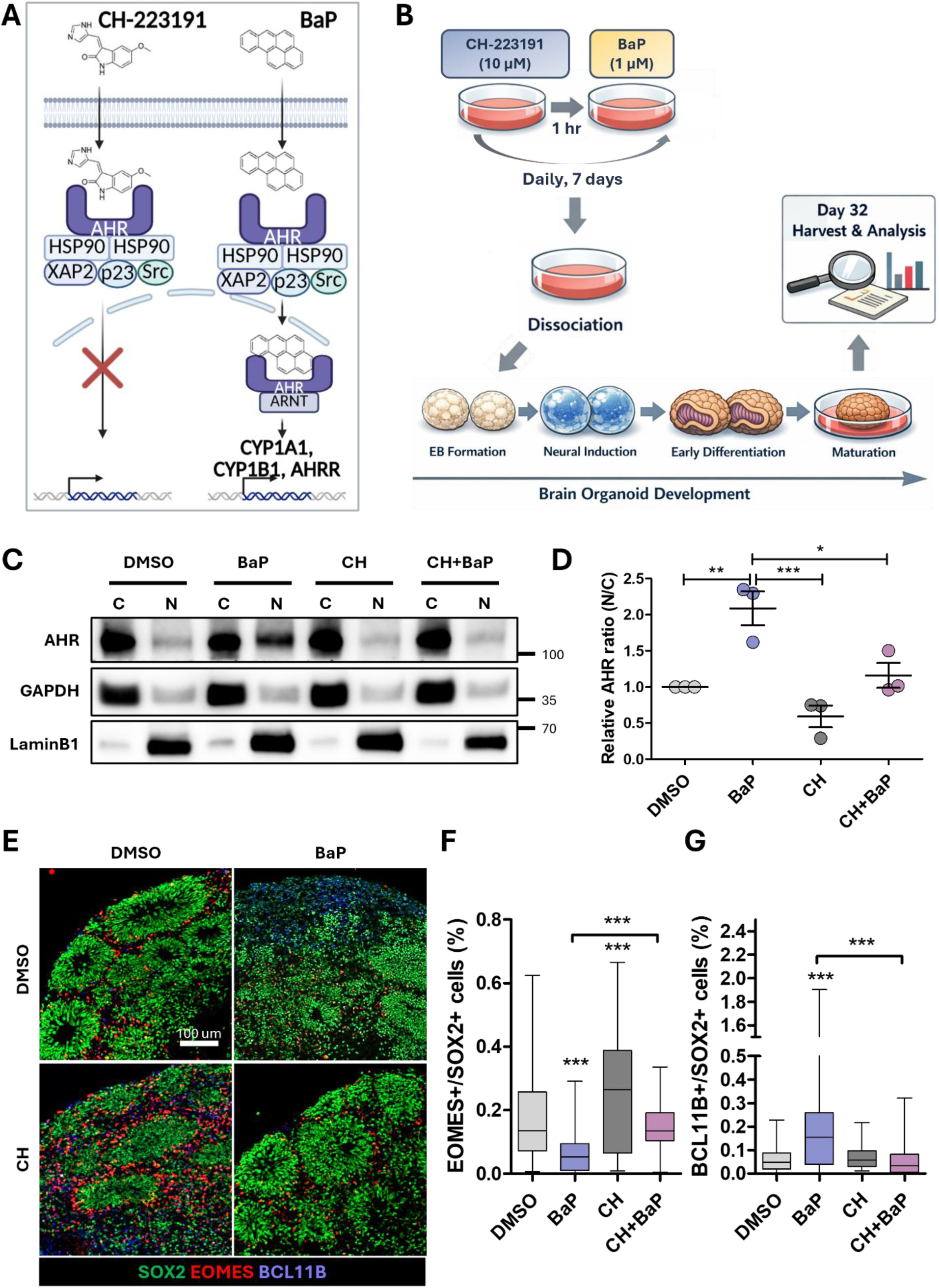
AHR antagonism by CH-223191 rescues BaP-induced alterations in AHR signaling and neurogenic differentiation. **(A)** Schematic illustration of BaP and CH-223191 (AHR antagonist) actions on AHR signaling. BaP activates AHR and promotes its nuclear translocation, whereas CH-223191 inhibits AHR translocation and activation. Created with BioRender.com. **(B)** Schematic diagram illustrating the rescue design. ESCs were pretreated with CH-223191 for 1 hr prior to BaP exposure, followed by organoid generation and downstream analyses. **(C** and **D)** Representative Western blot images and quantifications showing that BaP-induced AHR nuclear translocation is attenuated by CH-223191 pretreatment. **(E)** Representative immunofluorescence images of organoids stained for SOX2 (green), EOMES (red), and BCL11B (blue). **(F** and **G)** Box plots showing that the BaP-induced decrease in EOMES⁺ intermediate progenitor cells and increase in BCL11B⁺ neurons are restored by CH-223191 pretreatment. In this figure, CH indicates CH-223191 (10 µM) treatment and BaP indicates BaP (1 µM) treatment. Values represent mean ± SEM. *\*p < 0.05, **p < 0.005, ***p < 0.0005*. Scale bar: **(E)** 100 μm.

ESCs were pretreated with 10 µM CH-223191 for 1 h prior to BaP exposure, and this pretreatment paradigm was repeated daily during medium changes for 1 week (**Fig. 7B**). Following this exposure phase, ESCs were subjected to cerebral organoid differentiation, and day 32 organoids were analyzed to assess downstream neurodevelopmental outcomes. To first confirm effective inhibition of AHR activation at the ESC stage, cytosolic and nuclear fractionation was performed after 1 week of treatment, followed by Western blot analysis of AHR localization (**Fig. 7C**). BaP exposure markedly increased AHR nuclear translocation, as reflected by an elevated nuclear-to-cytosolic AHR ratio compared to DMSO controls (**Fig. 7D**). CH-223191 alone did not affect nuclear enrichment of AHR. Importantly, CH-223191 pretreatment significantly attenuated BaP-induced AHR nuclear accumulation, restoring the nuclear/cytosolic AHR ratio to levels comparable to controls. These findings confirm that BaP-driven AHR activation in ESCs is effectively suppressed by competitive antagonism.

We next examined whether the inhibition of AHR signaling could rescue the neurogenic alterations observed in BaP-derived organoids. BaP exposure significantly reduced the proportion of EOMES⁺ intermediate progenitor cells and increased the proportion of BCL11B⁺ neurons, consistent with premature neurogenic differentiation (**Fig. 7E-G**). Strikingly, CH-223191 pretreatment restored both EOMES⁺ and BCL11B⁺ cell populations to levels comparable to DMSO controls (**Fig. 7E-G**).

Collectively, these results demonstrate that BaP-induced AHR activation during the ESC stage is mechanistically linked to subsequent alterations in cortical lineage progression in cerebral organoids. Pharmacological inhibition of AHR signaling prevented both early molecular activation and later neurodevelopmental defects, suggesting that early AHR activation by BaP drives long-term neurodevelopmental abnormalities.

## 4. Discussion

This study identifies the pluripotent stage as a particularly sensitive developmental window for BaP exposure and suggests that transient molecular perturbation at this stage can influence later brain development. Although BaP exposure did not cause overt cytotoxicity or disrupt pluripotency in human ESCs, it was sufficient to activate AHR-associated transcriptional responses and subsequently alter lineage balance in derived cerebral organoids. The ability of AHR antagonism to reverse both early molecular responses and later developmental phenotypes further supports a mechanistic link between ESC-stage AHR activation and downstream neurodevelopmental consequences.

One notable aspect of the ESC response was the relatively modest degree of global transcriptional change despite clear pathway-level convergence on AHR-regulated processes. Rather than causing broad transcriptomic disruption, BaP appeared to engage a focused regulatory program centered on xenobiotic-responsive signaling. These findings suggest that early developmental toxicity may not require widespread gene expression changes; instead, selective pathway activation during pluripotency may be sufficient to bias later differentiation trajectories. Consistent with this idea, early transcriptional perturbations may remain phenotypically silent in stem cells, yet become amplified as lineage commitment proceeds [55–57]. Alternatively, or in parallel, AHR activation during the ESC stage may alter the epigenetic landscape during this critical developmental window. Changes in chromatin accessibility, histone modifications, or DNA methylation could persist beyond the initial exposure period and subsequently influence neural lineage progression and cortical development in organoids. In support of this possibility, KEGG enrichment of chemical carcinogenesis-DNA adduct pathways suggests that BaP metabolism in ESCs may generate reactive intermediates capable of covalently modifying DNA, a process previously linked to altered chromatin regulation and epigenetic instability [52, 58]. Interestingly, GO analysis also identified enrichment of piRNA processing pathways involving *PIWIL3*, *EXD1*, and *TDRD6*, genes associated with small RNA-mediated epigenetic regulation and transposon control during early development [59, 60].

AHR is expressed in blastocysts [61, 62]. *In vitro*, AHR expression was highest at the ESC stage and declined during early differentiation in both a prior study [21] and our model system. This temporal pattern suggests that ligand-dependent AHR activation may exert disproportionate influence on pluripotent stem cells, prior to the progressive restriction of developmental programs. In this context, AHR is likely functioning not only as a xenobiotic sensor but also as a developmental regulator whose activation alters the timing or balance of downstream lineage decisions [62]. The later organoid phenotype supports this interpretation. BaP exposure reduced neural progenitor populations while increasing early deep-layer neuronal output, indicating a shift away from progenitor maintenance toward neurogenic commitment. This type of early imbalance is consistent with mechanisms proposed to underlie several neurodevelopmental disorders in which altered progenitor dynamics precede broader structural and functional abnormalities of the brain [63–65]. Our findings therefore suggest that early transient AHR activation contributes to neurodevelopmental disorders associated with exposure to environmental pollutants by accelerating neural differentiation at the expense of neural progenitor maintenance.

The rescue experiments provide important mechanistic support for this interpretation. Pharmacological inhibition of AHR attenuated BaP-induced nuclear translocation in ESCs and normalized lineage proportions in derived organoids, indicating that AHR signaling contributes functionally to the observed phenotype rather than merely accompanying exposure. While BaP may engage additional pathways, the parallel recovery of early molecular and later developmental endpoints indicates that AHR-dependent signaling is a critical driver of BaP’s effects in this model. This is particularly relevant because the initial ESC phenotype was molecularly subtle, yet later developmental consequences were prominent, suggesting that early receptor-mediated signaling can leave persistent developmental effects even when immediate toxicity is minimal.The developmental stage modeled in this study suggests toxicological susceptibility of blastocysts. Human ESCs represent a pluripotent state corresponding to the inner cell mass of the blastocyst, approximating an early pre-implantation developmental window before establishment of a functional placenta or fetal circulation [66–68]. Although pre-implantation embryos are not directly connected to maternal blood, maternal exposure to small lipophilic compounds may still influence the embryo through the uterine environment before mature biological barriers are established [69–71]. This early developmental period has long been considered particularly vulnerable to environmental perturbation because rapid cell fate decisions occur in the absence of fully developed protective interfaces [72, 73]. Given its lipophilic nature, BaP can readily cross cellular membranes [7, 74], supporting the biological plausibility that even transient exposure during very early development may influence later developmental trajectories.

Although average daily human exposure via inhalation is generally lower than concentrations commonly used *in vitro* [8, 36, 75], exposure levels can be substantially elevated in populations residing near industrial emission sources in heavily polluted environments, where localized PAH burdens exceed background levels. In addition, occupational exposure in firefighters, military personnel, and workers in combustion-related settings may result in repeated high-intensity contact with PAH mixtures that cannot be overlooked [76–80]. BaP is highly lipophilic and readily partitions into biological membranes, allowing enrichment within tissues beyond what circulating concentrations alone may indicate [7, 35]. In addition, the physicochemical stability and hydrophobicity of PAHs contribute to their persistence across environmental compartments, including air, soil, dust, and contaminated food sources, which may increase cumulative human exposure under conditions of ongoing urban combustion and expanding wildfire events [7, 81]. For these reasons, low micromolar concentrations are widely used in mechanistic *in vitro* studies to approximate intracellular exposure levels sufficient to activate receptor mediated signaling pathways without inducing overt cytotoxicity [82, 83]. Under our experimental conditions, BaP exposure did not induce detectable ESC death, gross morphological disruption, or pronounced transcriptome changes, supporting the interpretation that the observed developmental effects primarily reflect signaling-mediated perturbation rather than nonspecific toxicity.

Several limitations should be acknowledged. Cerebral organoids capture major features of early corticogenesis but lack vasculature, immune interactions, and full systemic metabolism [84–86], all of which may influence toxicant responses *in vivo*. In addition, lineage changes were assessed at an early developmental stage, and longer-term consequences for neuronal maturation, circuit formation, or functional activity remain unresolved. Future studies incorporating single-cell transcriptomics, epigenomic profiling, and electrophysiological analyses will be important to determine how persistent these early BaP-induced alterations remain during later maturation.

In summary, our findings indicate that BaP exposure during pluripotency activates AHR signaling and induces subtle early molecular changes that are sufficient to alter subsequent cortical lineage progression in human cerebral organoids. By demonstrating that pharmacological inhibition of AHR prevents these effects, this study identifies AHR-mediated signaling as a mechanistic link between early PAH exposure and later neurodevelopmental imbalance.

## 5. Abbreviations

AHR: Aryl hydrocarbon receptor
AHRR: Aryl hydrocarbon receptor repressor
ARNT: Aryl hydrocarbon receptor nuclear translocator
BaP: Benzo[a]pyrene
CYP: Cytochrome P450
DEG(s): Differentially expressed gene(s)
EB: Embryoid body
ED: Early differentiation
ESC(s): Embryonic stem cell(s)
GO: Gene ontology
GSEA: Gene set enrichment analysis
KEGG: Kyoto encyclopedia of genes and genomes
LDH: Lactate dehydrogenase
Mat: Maturation
NI: Neural induction
PAH(s): Polycyclic aromatic hydrocarbon(s)
PPI: Protein-protein interaction

## 6. CRediT authorship contribution statement

**Bohyeon Jeong**: Conceptualization, Methodology, Investigation, Data curation, Visualization, Funding acquisition, Writing - original draft, review & editing. **Lei Yang**: Methodology, Investigation, Formal analysis. **Tharindu Ranathunge**: Methodology, Investigation, Formal analysis. **Young-Goo Han**: Supervision, Resources, Funding acquisition, Writing - review & editing.

## 8. Declaration of generative AI and AI associated technologies in the manuscript preparation process

During the preparation of this work, the authors used an AI-assisted image generation tool to support the design of the experimental timeline illustration. The authors reviewed, edited, and finalized the figure and took full responsibility for the content of the published article.

## 9. Acknowledgements

This study was supported by the National Research Foundation (NRF) of Korea (RS-2024-00406672 to B.J.), the American Lebanese Syrian Associated Charities (ALSAC), and the National Institute of Neurological Disorders and Stroke, National Institutes of Health (R01NS100939 to Y.-G.H). The content is solely the responsibility of the authors and does not necessarily represent the official views of the National Institute of Health.

We are grateful to Lei Li and Hongjian Jin from the Center for Applied Bioinformatics at St. Jude Children’s Research Hospital for their assistance with sequencing analysis and bioinformatics support. We also appreciate the support of the Developmental Neurobiology Imaging Center at St. Jude Children’s Research Hospital for providing microscopy support and access to imaging facilities. We thank the Demontis Lab for access to the ChemiDoc imaging system, and Dong Geun Lee for introducing us to colleagues in the Department of Chemical Biology and Therapeutics, which enabled the analysis of intracellular BaP accumulation. We further thank Kris M. Olesen for valuable feedback on the manuscript, as well as members of the Han Lab for helpful discussions.

